# Towards establishing a 24-hour, microplate-based, transcriptomics assay for rainbow trout embryos

**DOI:** 10.1101/2023.07.07.547968

**Authors:** Niladri Basu, Aylish Marshall, Hugo Marchand, Emily Boulanger, Krittika Mittal, Jessica Head

## Abstract

There is interest in the development of early-life stage (ELS) tests with fish embryo models that are high-throughput and can generate transcriptomics point of departure (tPOD) values. The objective of this study was to establish a method in rainbow trout (*Oncorhynchus mykiss*) hatchlings that could satisfy both of these interests. We based our pilot method on recent efforts by U.S. EPA researchers to establish a larval fathead minnow high throughput transcriptomics assay. Here, 1-2 day post hatch trout were assayed in 24-well plates in which they were exposed for 24 hours to 12 different concentrations of test chemicals, including a negative control (DMSO, culture water). Test concentrations were anchored with a chemical’s LC50 data from the US EPA ECOTOX database and EnviroTox database, and from this, concentrations were spaced on a half-log basis that spanned 6-7 orders of magnitude. In pilot study 1 we tested 3,4-dichloroaniline, CuSO_4_ (0.34 mg/L), and ethinylestradiol. In pilot study 2 we tested 3,4-dichloroaniline (58.5 mg/L), CuSO_4_ (0.34 and 0.41 mg/L), ethinylestradiol (>10 µg/L), permethrin (>10 µg/L), malathion (0.61 mg/L), 6PPD quinone (5.6 µg/L), acetaldehyde (41.2 mg/L), 4-fluoroaniline (242.7 mg/L), glyphosate (∼150 mg/L), ethanol (>1 g/L), thiamethoxam (>300 mg/L), and allyl alcohol (>30 mg/L). In both pilot studies derived LC50 values are provided in parentheses. Repeated studies of CuSO_4_ yielded consistent LC50 values (0.34, 0.34, 0.41 mg/L). The correlation between LC50s from the current study for rainbow trout embryos versus those from the literature on adult rainbow trout for 7 chemicals was r^2^ = 0.91. Work is underway to optimize transcriptomics assays from these samples using EcoToxChips and UPXome, with the ultimate goal to be able to derive transcriptomics points of departure. Taken together these results provide a foundation towards establishing a novel testing platform for chemical and environmental risk assessment that is much quicker (24 hrs), ethical (non-protected life stages), resource efficient (e.g., microplate-based, small volumes of chemicals), and more informative (molecular clues into MOA) than traditional bioassay approaches.

## 1. INTRODUCTION

Regulatory frameworks for assessing chemicals are highly dependent on data from live animal-toxicity tests that follow standard protocols. Fish are one of three taxonomic groups used to assess chemical hazards to aquatic ecosystem health (United Nations, 2015). In general, fish toxicity tests involve exposing organisms to a graded range of chemical concentrations, and observing impacts on growth, reproduction and survival. From the resulting dose-response data, benchmark doses (BMD) are calculated and used to estimate predicted no-effect concentrations (PNEC) deemed to be “safe”. Such “kill ‘em and count ‘em” methods are highly tangible, institutionalized, and provide direct evidence of the concentration of a chemical that causes harm, but they have shortcomings that represent major barriers. For one, these tests are prohibitively expensive and require numerous animal lives; we estimated that standardized tests involving fish may cost between $16,000 and $412,000 USD, use about 150 animals, and take two months (Mittal et al., 2022). Given that several of these tests are typically required to render a decision on a chemical’s safety, testing a single chemical may take up to several years and cost millions of dollars. Furthermore, even when data from a full suite of tests are available, they only report on a narrow set of toxicological endpoints. These cannot capture the full range of biological impacts or provide information on the mechanism of toxicity. Thus, regulatory agencies and industry are moving toward New Approach Methods (NAMs) to support risk assessments to meet new legislative mandates, while reducing animal use, costs, and time required for testing (EChA 2016; US EPA, 2018; van der Zalm et al., 2022).

The BMD concept is now being applied to molecular data (National Toxicology Program 2018). Prior to adverse outcomes at the organismal level, molecular responses such as changes in gene expression (transcription) occur. The concentration resulting in a concerted transcriptomic change indicates a transcriptomic point of departure (POD) (Johnson et al., 2022). There are a growing number of studies documenting that tPODs derived from short-term studies yield values that are lower in concentration than BMDs derived from chronic animal bioassays that focus on adverse apical outcomes. This concept is now being extended to studies on fish. For example, in an analysis of 5 transcriptomics datasets concerning fish exposed to endocrine disrupting compounds (bisphenol A, ethinylestradiol, and diethylstilbestrol), Pagé-Larivière et al. (2019) calculated that the tPOD values were within one order of magnitude of lowest observable effect concentrations (LOECs) determined from chronic fish bioassays.

There is now interest in the development of methods with high-throughput potential that couple early-life stage (ELS) tests with non-protected fish embryo models (as an alternative to animal test methods) with transcriptomics approaches that may lend themselves to the derivation of tPOD values. Towards this, U.S. EPA researchers have recently proposed, and pilot tested such a method with larval fathead minnow (Villeneuve et al., 2022). Specifically, hatchlings were exposed in microplates to ten test chemicals (at 12 concentrations spanning six orders of magnitude) for 24 hrs following which the whole transcriptome was sequenced and analyzed to derive tPOD values. In another case, researchers from Canada’s National Research Council (NRC) have refined standard test methods on zebrafish to derive tPOD values and relate them with several phenotypic measures (Morash et al., 2023).

Rainbow trout are another fish species for which a high-throughput, ELS-based toxicogenomic test method may prove helpful. Trout are the main regulatory fish model used in Canada (i.e., >84,000 individuals/yr used through test EPS1/RM/13 OR ECCC EPS1/RM/28 largely for compliance monitoring of municipal waterways and effluents), and are also used across the European Union. Accordingly, building upon recent efforts in fathead minnow and zebrafish, the objective of the current study was to establish an ELS-based, toxicogenomic test method for rainbow trout that has the potential to be high-throughput. Secondary objectives included to: a) pilot test this method on chemicals with diverse properties; b) compare the resulting data from this method against information on adults; and c) compare the repeatability of the test. As articulated in the Methods section, much of the test design was motivated by recent efforts from Dr. Dan Villeneuve’s group at the US EPA focused on similar objectives for fathead minnows.

## 2.0 METHOD

### 2.1 Assay design considerations

The design of this ELS-based, toxicogenomic test method for rainbow trout was motivated by an effort by U.S. EPA researchers to establish a similar method in larval fathead minnows (Villeneuve et al., 2022). In particular, we draw from a sentiment by Villeneuve et al. (2022) that there is likely no single assay design that will serve the universe of all chemicals (e.g., vary in ADME properties, mechanism of action), and so practical and pragmatic reasons along with pertinent biological considerations should feature prominently towards assay design and standardization.

2.1.1 Life stage. Similar to the work by Villeneuve et al. (2022) on larval fathead minnows, we focused on rainbow trout that were ∼1 day post hatch. At this stage, trout hatchlings (eleutheroembryos i.e., ‘free embryo’) are considered a non-protected species as they are not free feeding i.e., they depend on their yolk sac for nutrition (Belanger et al., 2010; Halder et al., 2010) (Figure 1). A focus on this life stage also negates the need to introduce food into the test system, which may raise concerns over water quality and interaction between feed components and test chemicals.

**Figure 1.**
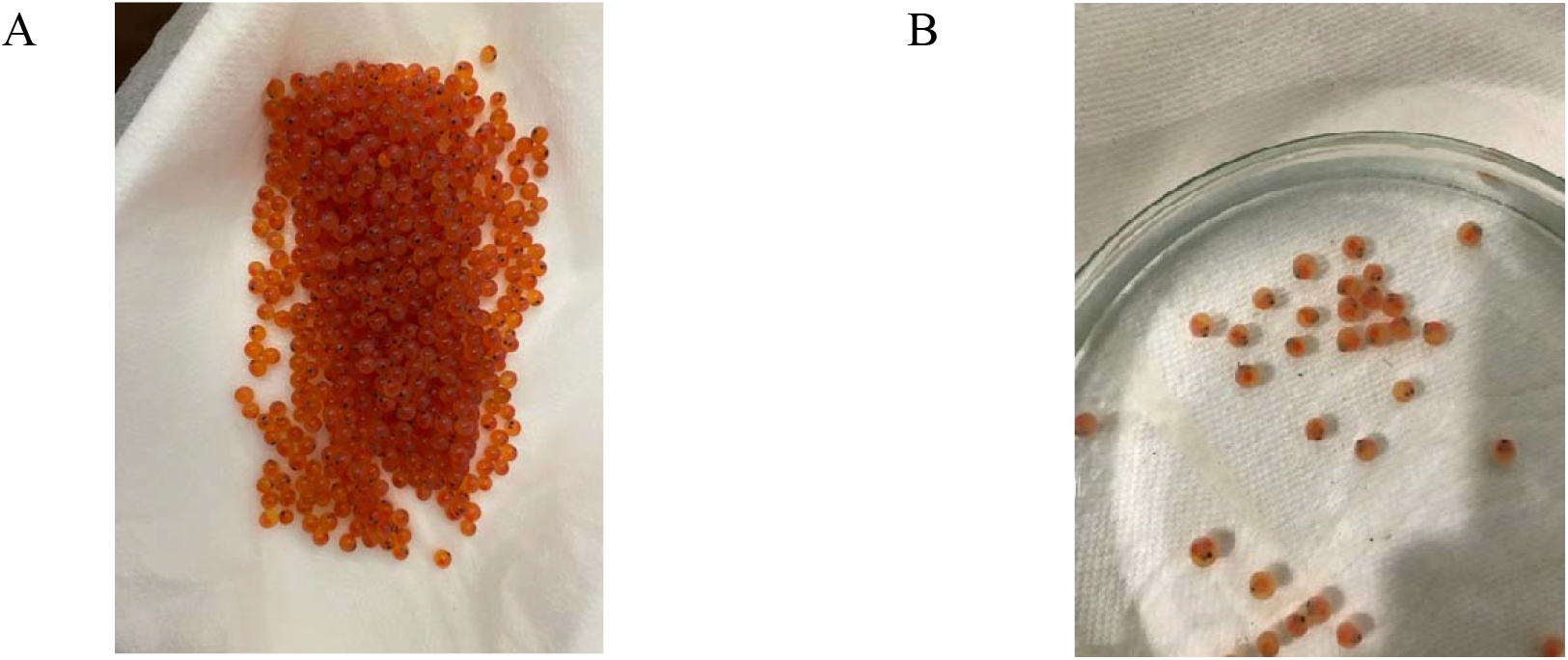
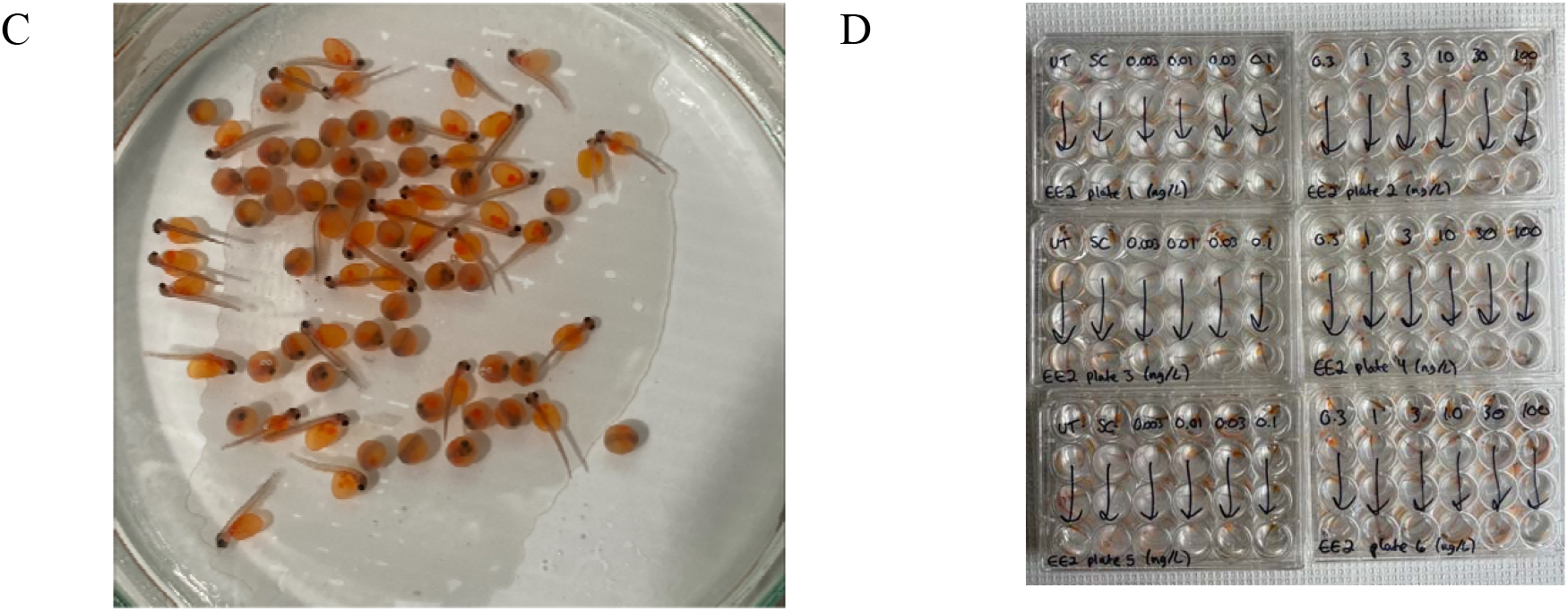
Photographs of trout embryos at: A) receipt from transportation; B) as eyed eggs during initial inspections; C) as eleutheroembryos; and D) in exposure plates.

There are also biological considerations associated with exposures during the eleutheroembryo phase. Belanger et al. (2010) argued that this phase is one of the most vulnerable in terms of sensitivity to chemical or physical insults. Even though fish at this life stage are not fully developed, organs have been formed and transcriptomic changes associated with early development have likely finished. Finally, the hatchlings are mobile and responsive to stimuli thus permitting behavioral observations.

2.1.2 Chemical treatment. Following the approach of Villeneuve et al. (2022), each test chemical will be tested at 11 concentrations as well as a water or solvent control (=12 test concentrations in total). The highest test concentration was the reported LC50 value obtained from the US EPA ECOTOX or EnviroTox databases (Table 1). From these databases, we prioritized data from rainbow trout that underwent 24 – 96 hour waterborne exposures. From this highest concentration, lower concentrations were determined on a half-log scale. For example, if the highest test concentration was 320 ug/L, then the subsequent concentrations would be 100, 32, 10, 3.2, 1, 0.32, 0.1, 0.032, 0.01, 0.0032, and 0 ug/L [=12 test concentrations, spread over 5 orders of magnitude]. Note that this was the exposure design in pilot #1, and that the design changed for pilot #2 as discussed further below based on lessons learned.

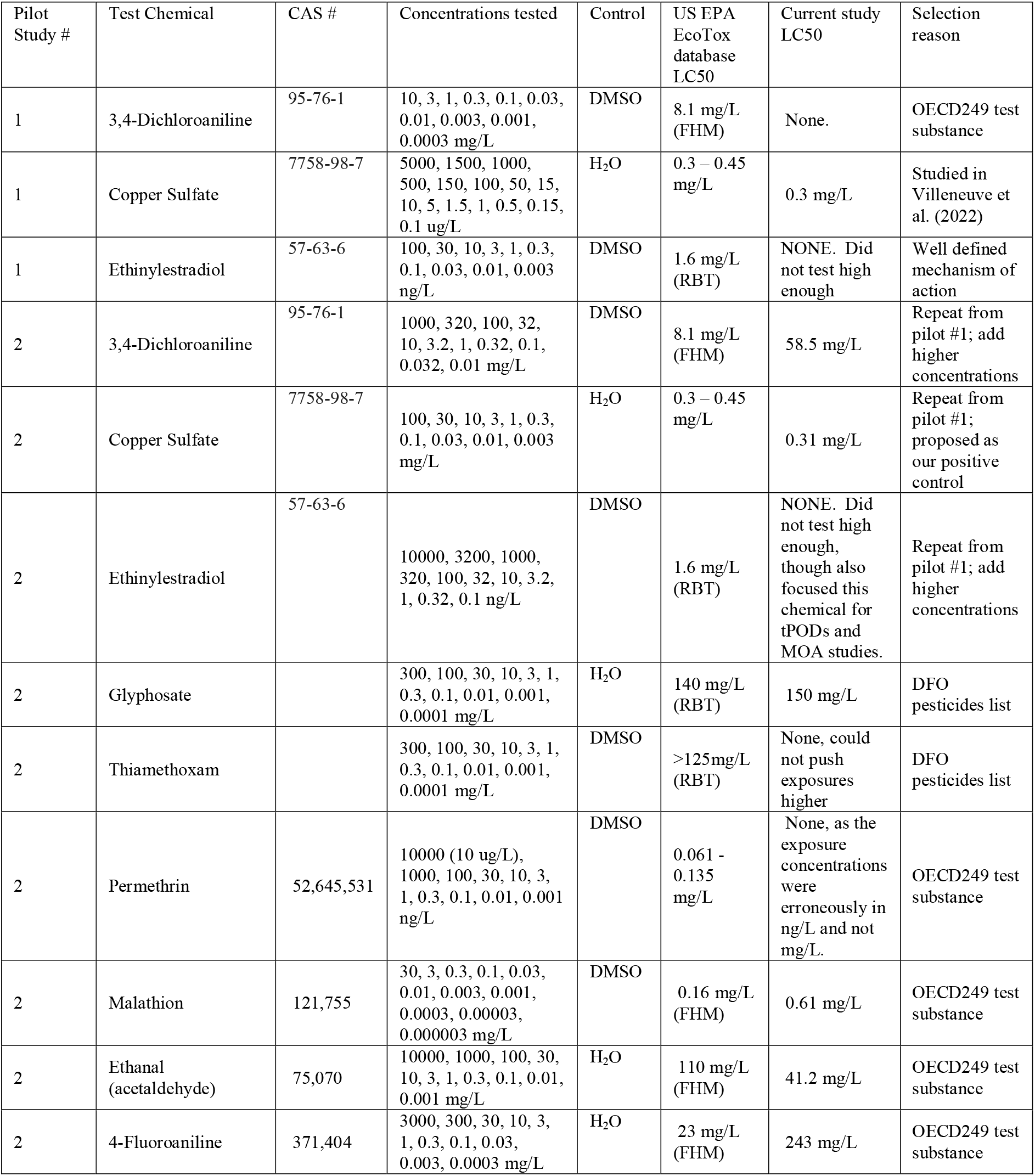
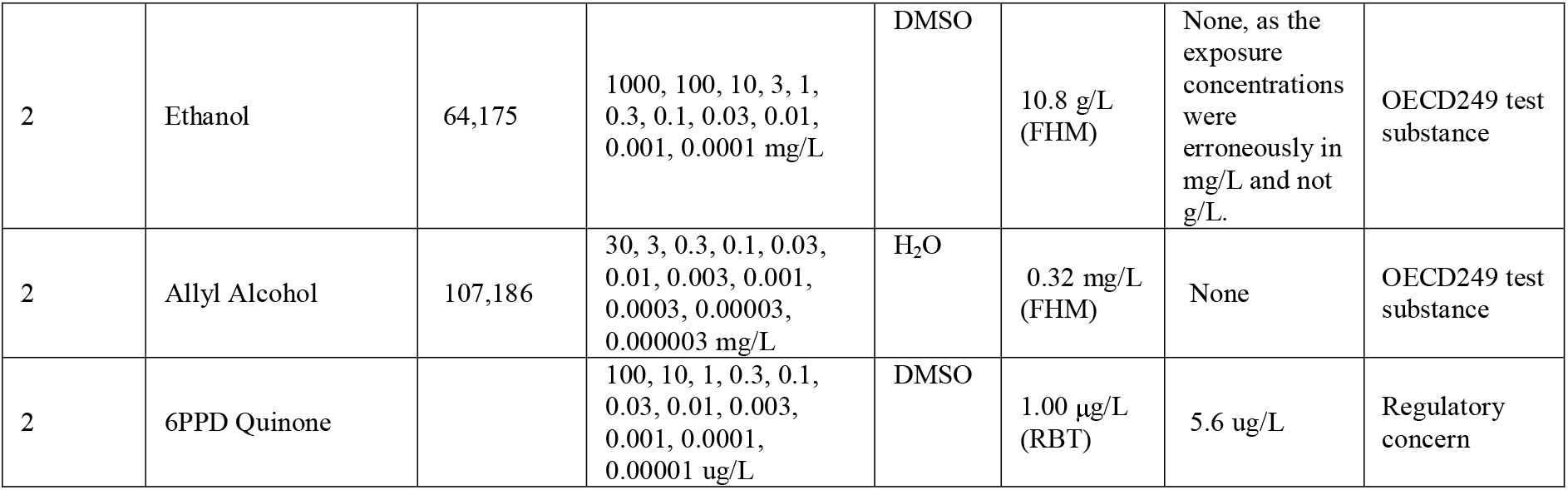

In terms of negative controls, we favored the use of DMSO or water as a solvent. In the current study, the maximal DMSO concentration was set at 0.5% similar to Villeneuve et al. (2022). A previous study on zebrafish embryos concluded that DMSO up to 1% is acceptable (Hoyberghs et al., 2021). In a future study, we will assess molecular changes associated with a range of DMSO exposures in this test system.

In a typical study we expect that there is only one biological replicate given that we source the trout eggs from a single hatchery at a given time. In terms of technical replicates, this was determined from the following considerations: a) RNA yield per embryo and downstream transcriptomic analysis needs; b) number of test concentrations to yield a tPOD; c) replicates of a given test concentrations; d) total number of microplate wells required; and e) exposure layout across the microplates. The base method involves 12 embryos per test concentration, and splitting these into 4 pools of 3 or 4 embryos (i.e., 4 technical replicates with 4 pooled individuals per replicate in pilot #1 and 3 pooled individuals in pilot #2). This design adhered to the recommendations from Villeneuve et al. (2022).

2.1.3 Exposure duration. Similar to the work by Villeneuve et al. (2022), we decided upon a single 24-hour exposure. This was likely enough time to elicit a measurable transcriptomic response and not raise water quality concerns (Flynn et al. unpublished observations in Villeneuve et al., 2022). It would also minimize a need to replenish test solutions which could introduce additional errors in the experiment (e.g., pipetting) or stress to the organism. Finally, a static 24-hour exposure period aligns with our overall goal to realize an efficient method.

2.1.4 Exposure chambers. To align with broader efforts to realize test systems with throughput potential (i.e., screen tens to hundreds of chemicals/samples in a week), designing the assay within microplates was desired. Owing to the larger size of the trout eggs and eleutheroembryos versus fathead minnows/zebrafish (Figure 1), the assay was developed to function in 24-well plates (∼15mm diameter, 17 mm depth). The well volume in such plates is 3.4 ml and thus ∼10-fold higher than the volume of a typical 96-well microplate. We set an initial target of 2 mL of test solution per well. There are unpublished observations by Flynn et al. (in Villeneuve et al., 2022) that 0.3 ml was sufficient to maintain water quality over a 24 hour static exposure scenario with larval fathead minnow. The larger volume here would provide extra buffering against possible impairments to water quality from chemical exposure (e.g., pH changes, ammonia levels). We also expect 2 ml to provide a larger exposure volume for the eleutheroembryo trout as well as ample water for the organism to move freely.

### 2.2 Rainbow trout

All female diploid rainbow trout eggs at the eyed-stage were obtained from Troutlodge (Bonney Lake, WA, USA). They were air transported to McGill University in damp paper towels in an insulated container with ice so that the transport temperature likely ranged from 4 to 12°C. Required import permits from the Canadian Food Inspection Agency (CFIA) were obtained.

Once at McGill University, the eggs were split into medium-sized beakers (∼1L) filled with 12°C reconstituted hard water (ultrapure water supplemented with sodium bicarbonate [96 mg/L], magnesium sulfate [60 mg/L], calcium chloride [39 mg/L], and potassium chloride [4 mg/L]; modified US Environmental Protection Agency, USEPA, 2002). After 30 minutes of acclimation, the eggs were poured into a large glass petri dish to identify and remove dead ones. Next, about 40 - 60 eggs and water from the petri dish were moved into 200mL air-powered tumblers (Ziss ZET-E55 fish incubator), which were then submerged into 40 L glass aquaria upon which the tumblers were connected to an air source. Air valves were open just enough to provide gentle movement of eggs. About 24 – 48 hours prior to receiving eggs, the glass aquaria were filled with reconstituted hard water and allowed to acclimate in a walk-in cold room with an air temperature set to 12 ± 1°C (Berube et al., 2022). Eggs were kept in total darkness and only exposed to dimmed light to allow for daily inspection and removal of dead eggs (Di Lombo et al., 2021; Weeks Santos et al., 2021).

In the current work we conducted two pilot studies. For pilot study #1 the eggs were received on January 26, 2023, with hatching commencing on February 1, 2023. The exposures were initiated on February 2, 2023 between 11am and 2:55pm, and they ended 24 hours later. For pilot study #2 the eggs were received on March 30, 2023, with hatching commencing on April 3, 2023. The exposures for some chemicals were initiated on April 4, 2023, and on April 5 for some other chemicals.

### 2.3 Chemical Solutions

The test chemicals are detailed in Table 1.

In pilot #1, each test chemical was tested at 12-15 concentrations including a negative control (Table 1). The highest test concentration was the reported LC50 value obtained from the US EPA EcoTox database.

In pilot #2, a tapered exposure design was pursued based on the results from pilot #1:

- Anchor concentration: The test chemical’s reported LC50 value was obtained from the US EPA EcoTox database, and this value was set as the third highest test concentration.
- LC50 push concentrations: The two highest test concentrations were 10-fold and 100-fold higher than the anchor concentration to push the test system in a zone of adversity/toxicity, and potentially help calculate a crude LC50 value.
- tPOD zone: The middle six concentrations were assigned to span a semi-log scale. For example, if the highest test concentrations are 1000 and 100 ug/L (LC50 push values), then the subsequent concentrations will be 10 (i.e., LC50 value from US EPA EcoTox database), followed by 3.2, 1, 0.32, 0.1, 0.032, and 0.01 ug/L.
- Zero zone (n = 3 concentrations): To ensure hitting a baseline (null) gene expression range, the bottom 2 concentrations will be 10x and 100x lower (inverse of LC50 push range). If the last TPOD range value is 0.032 ug/L, we would have 0.0032, and 0.00032 ug/L, with a final value of 0 for the negative control. For pilot 2, each chemical was tested against 12 concentrations spread over 7.5 orders of magnitude. In pilot 2, a decision was also taken for all chemicals to follow a log10 scale that included half-log values of 3.2. Thus, if the anchor concentration yielded a value of 7 ug/L, we rounded that up to 10 ug/L.

### 2.4 Exposures and termination

About 5 – 7 days after receiving eggs, the first signs of hatching were observed. We aimed to discard the first 5% of early hatchlings (as well as 5% of last hatchlings) to avoid potential outlier effects. During inspection in the petri dishes, eleutheroembryos were removed one at a time with a turkey baster and placed into an empty well in the 24 well microplates (Fisherbrand™ FB012929) along with ∼2-3 mL of water. Once 6 plates (i.e., 144 wells) were filled completely with eleutheroembryos, then a chemical exposure commenced. Working in teams, the water was aspirated and quickly replaced with 2 mL of the pre-cooled exposure solution. All plates were pre-labelled, and following addition of exposure solutions they were covered with a lid and kept in shelves within the walk-in cold room. Exposure media at time zero was collected and stored frozen for three treatments (the database LC50 exposure group and two lower exposure groups). It took approximately 15 minutes to perform one chemical exposure.

Following a 24-hour exposure period, the plates were removed from the cold room and observations of the exposed eleutheroembryos noted. Foremost was numeration of any dead embryos. All plates were also photographed. Using a transfer pipette, the eleutheroembryos were removed and placed into a 2 ml tube for downstream transcriptomic analysis. Three to four eleutheroembryos from a given treatment group were pooled into one tube, and the tube was immediately frozen on dry ice. This resulted in 4 pools of 3 or 4 individuals per concentration (assuming no mortality). All samples were then moved into a designated box and stored in a – 80°C freezer.

Exposure media at termination was collected and stored frozen for the same three treatments that were sampled at time zero.

### 2.5 Data Analysis

Mortality data was tabulated and summarized. LC50 were calculated using the drc package in R V4.2.2.

## 3.0 RESULTS AND DISCUSSION

### 3.1 Mortality and hatching rates

Mortality and hatching rate are important characteristics of organismal health and study quality. Across both studies the overall hatching rate of rainbow trout was ∼97% (Table 2). These data align with OECD 236 (FET Zebrafish) in which the validity of the test calls for hatching rate to be >80% by the end of the 96-hour exposure period.

**Table 2.** Mortality and hatching rates of rainbow trout eggs and embryos.

|  | # Eggs received | Transport (# of mortalities, %) | Rearing and Hatching (# of mortalities, %) | Hatching Rate (#hatched, %) |
| --- | --- | --- | --- | --- |
| Pilot study #1 | 778 | 3 (0.4%) | 17 (2.2%) | 758 (97.8%) |
| Pilot study #2 | 2,200 | 0 (0%) | 48 (2.2%) | 2,152 (97.8%) |
| Total | 2,978 | 3 (0.1%) | 65 (2.2%) | 2,910 (97.8%) |

Upon commencement of exposures the overall mortality in the control plates was 4.4% (Table 3). These data align with OECD 236 (FET Zebrafish) in which the validity of the test calls for embryonic survival to be >90% in the negative controls (water or solvent) during the course of the 96 hour exposure period.

**Table 3.** Mortality in control wells.

|  | DMSO Controls (# of mortalities/# of hatchlings tested) | RHW Controls (# of mortalities/# of hatchlings tested) | Total (# of mortalities/# of hatchlings tested) |
| --- | --- | --- | --- |
| Pilot study #1 | 0/24 | 0/60 | 0/84 (0%) |
| Pilot study #2 | 4/60 | 6/84 | 10/144 (6.9%) |
| Total | 4/84 (4.8%) | 6/144 (4.2%) | 10/228 (4.4%) |

### 3.2 Repeatability

A key characteristic of study quality is assay repeatability. In both pilot studies, CuSO_4_ was included, and in pilot #2 the chemical was tested twice (i.e., using early-to-mid hatchlings and late hatchlings) to see if there may be differences due to developmental stage (Figure 2). The LC20 and LC50 values in all three cases were close. In pilot #1 the LC50 values for CuSO_4_ were 0.34 (0.33-0.35) and 0.41 (0.40-0.41), respectively (lower and upper standard error in parenthesis). In pilot #2, the LC20 and LC50 values were 0.29 (0.29-0.29) and 0.31 (0.31-0.31), respectively, among early to mid-hatchlings. Among the late hatchlings, the LC20 and LC50 values were 0.29 (0.29-0.30) and 0.33 (0.32-0.33), respectively. In all cases there was no mortality in the 0.1 mg/L CuSO_4_ group and 100% mortality in the 1 mg/L exposure groups.

**Figure 2.**
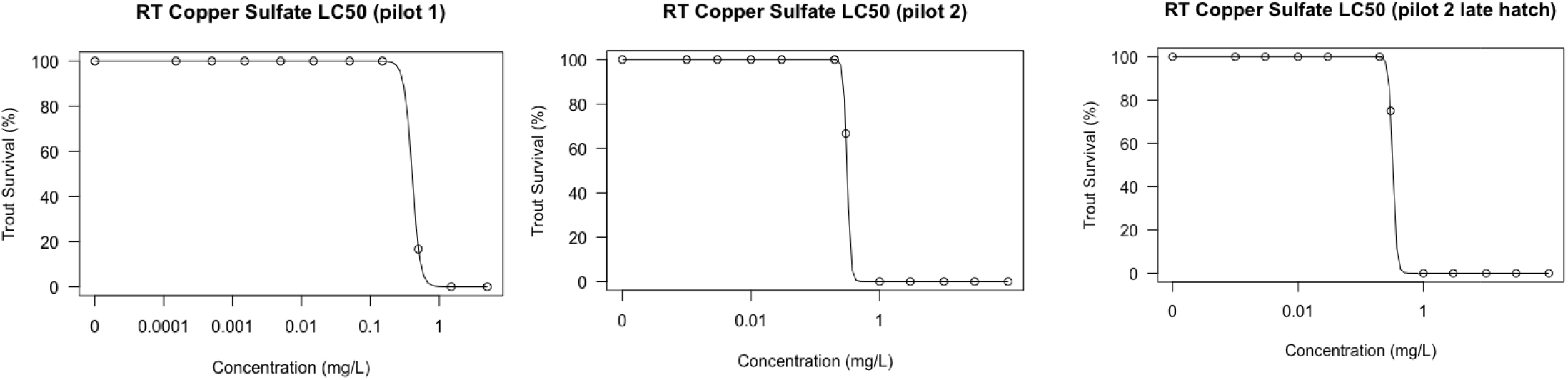
Survival plots for rainbow trout embryos exposed to CuSO_4_.

We also note that 3,4-Dichloroaniline and Ethinylestradiol were tested in both pilot studies. In pilot #1, no mortality suggestive of a LC50 (or even LC20) was observed. In an effort to derive LC values for both chemicals, in pilot #2 their maximum test concentrations were increased from 10 mg/L to 1,000 mg/L for 3,4-Dichloroaniline, and 100 ng/L to 10,000 ng/L for Ethinylestradiol. In doing so, a LC50 was realized for 3,4-Dichloroaniline (i.e., 58.5 mg/L) but not for Ethinylestradiol.

OECD 236 (FET Zebrafish) calls for a minimal mortality of 30% following exposure of zebrafish for 96 hrs to 4.0 mg/L 3,4-dichloroaniline. Our data from both pilot studies show that such a concentration of 3,4-Dichloroaniline is not toxic to rainbow trout. Rather, a concentration about 10-fold higher (e.g., 40 mg/L) would be expected to cause 30% mortality.

### 3.3 Comparisons with Literature

Across the two pilot studies, 12 chemicals were tested. Of these, we calculated LC50s for six chemicals and for another (glyphosate) we “eyeballed” a LC50 of 150mg/L (Table 1; Appendix 1). Of the remaining chemicals, for four of them (permethrin, ethinylestradiol, thiamethoxam, ethanol) we could not test higher concentrations (i.e., ones that would be ∼100-fold higher than the literature LC50) owing to concerns over solubility and/or procurement cost. The highest concentration of allyl alcohol that was tested was ∼100-fold greater than the reported LC50 though we did not find any adverse outcomes here.

Focusing on the seven chemicals for which LC50s were derived, we see good concordance between our data and the LC50s reported in the literature (Figure 3).

**Figure 3.**
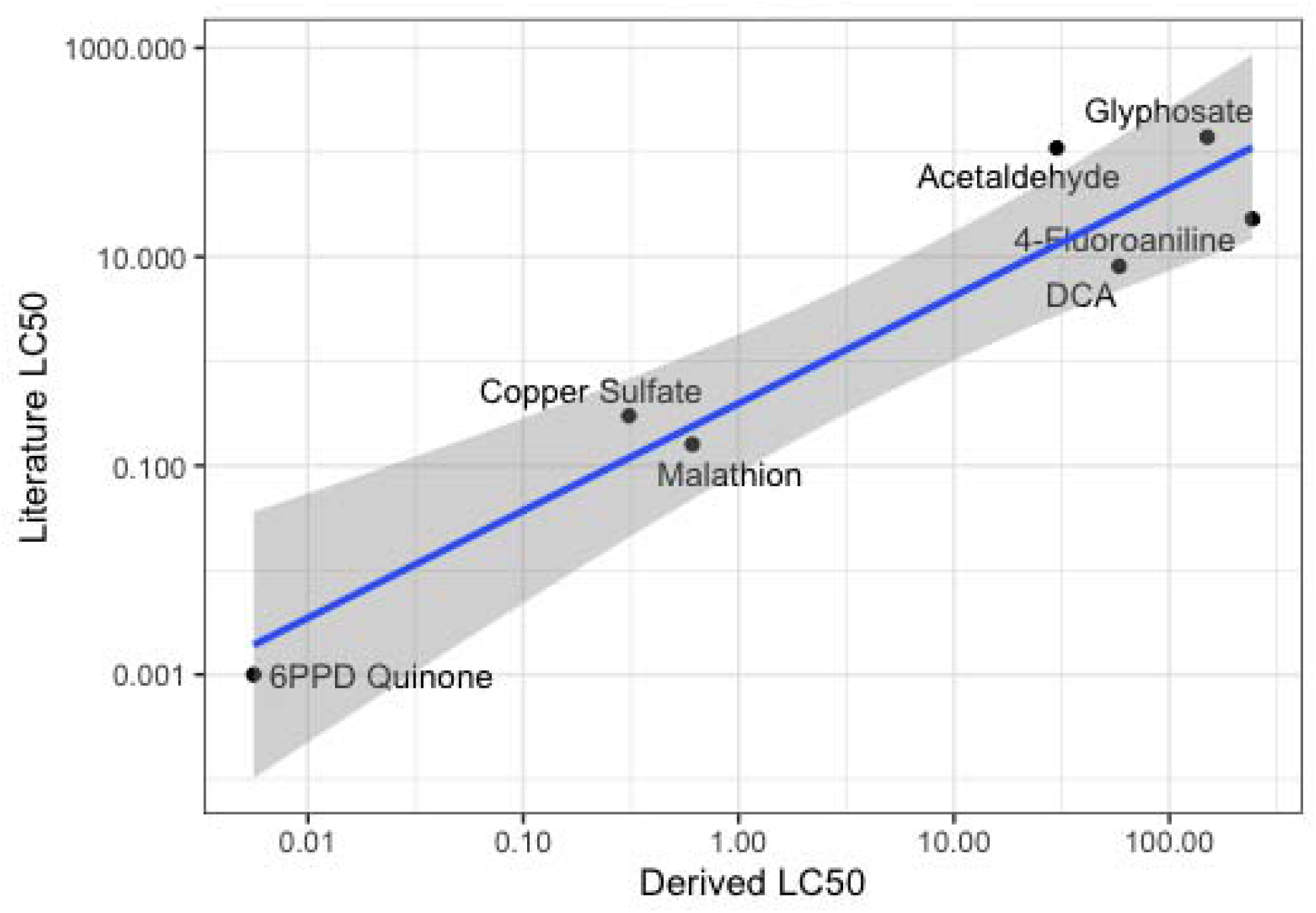
Comparison of LC50s from the current study for rainbow trout embryos (x-axis) versus those from the literature on adult rainbow trout (y-axis). The line of best fit is represented as y = 0.88x + 0.42, r^2^ = 0.91. We also note that the derived LC50 for glyphosate was estimated based on a review of the data (see Appendix) while for all other chemicals this value was calculated with the drc package in R.

There is considerable regulatory and ecological concern over 6PPD Quinone (Chen et al., 2023). Brinkmann et al. (2022) calculated a LC50 of 1 ug/L in ∼2 year old rainbow trout exposed for 72 hrs, and a LC50 of 0.59 in ∼1 year old brook trout exposed for 24 hrs. Di et al. (2022) calculated a LC50 of 1.7 – 4.3 ug/L of rainbow trout (∼3 g) exposed for 96 hours. In the current study we calculated a LC50 of 5.6 ug/L, which aligns well with previous studies on rainbow trout. Moreover, our study was based on an early lifestage model and was more efficient in several regards (e.g., microplate design with small test volumes, 12 concentrations tested, 24 hrs).

### 3.4 Transcriptomics Research

A key goal here was to develop a method that could relatively efficiently produce LC50 data as well as yield tissue for transcriptomics work. Our initial studies indicate that individual rainbow trout hatchlings (∼1-2 days post hatch) are ∼30 mg in size, and yield about 0.5–1.5 µg RNA. This is sufficient RNA for most transcriptomic applications (e.g, EcoToxChips require 1 ug, and RNAseq libraries require <100 ng).

Our proposed plan will be to pool 3 embryos and thus realize n=3 or 4 pools per test concentration. In doing so, a study of 29 pools of 3 embryos each yielded 30 ul of elution volume with total RNA of 2.7 ug/ml (SD: 0.16) with a minimum value of 2.4 and a maximum value over 3.0.

### 3.5 Concluding Remarks

Based on our review of the overall results, we contend that this test system has potential to serve as an ELS-based, toxicogenomic test method for rainbow trout that has the potential to be high-throughput. The results suggest that this method can work against a range of chemicals with diverse properties, and that for many substances that there is a strong concordance between the LC50s derived from this method and those in the literature based on whole animal studies. We also demonstrate good assay repeatability.

## Acknowledgements

We thank Drs. Peter Hodson and Julia Adams from Queen’s University and Roxanne Bérubé From Institut national de la recherche scientifique (INRS) for discussions on trout bioassays, as well as Dr. Dan Villeneuve and Kevin Flynn from the U.S. Environmental Protection Agency for several exchanges on their efforts to establish a high-throughput toxicogenomic method for larval fathead minnow. We thank Jenny Eng for administrative support. This project is part of a Genome Canada GAPP project entitled “Validation of the use of the EcoToxChip test system for regulatory decision-making”, and we thank the following financial sponsors of this work: Genome Canada, Génome Québec, and Environment and Climate Change Canada.

## Appendix 1 LC50 Figures of Test Chemicals

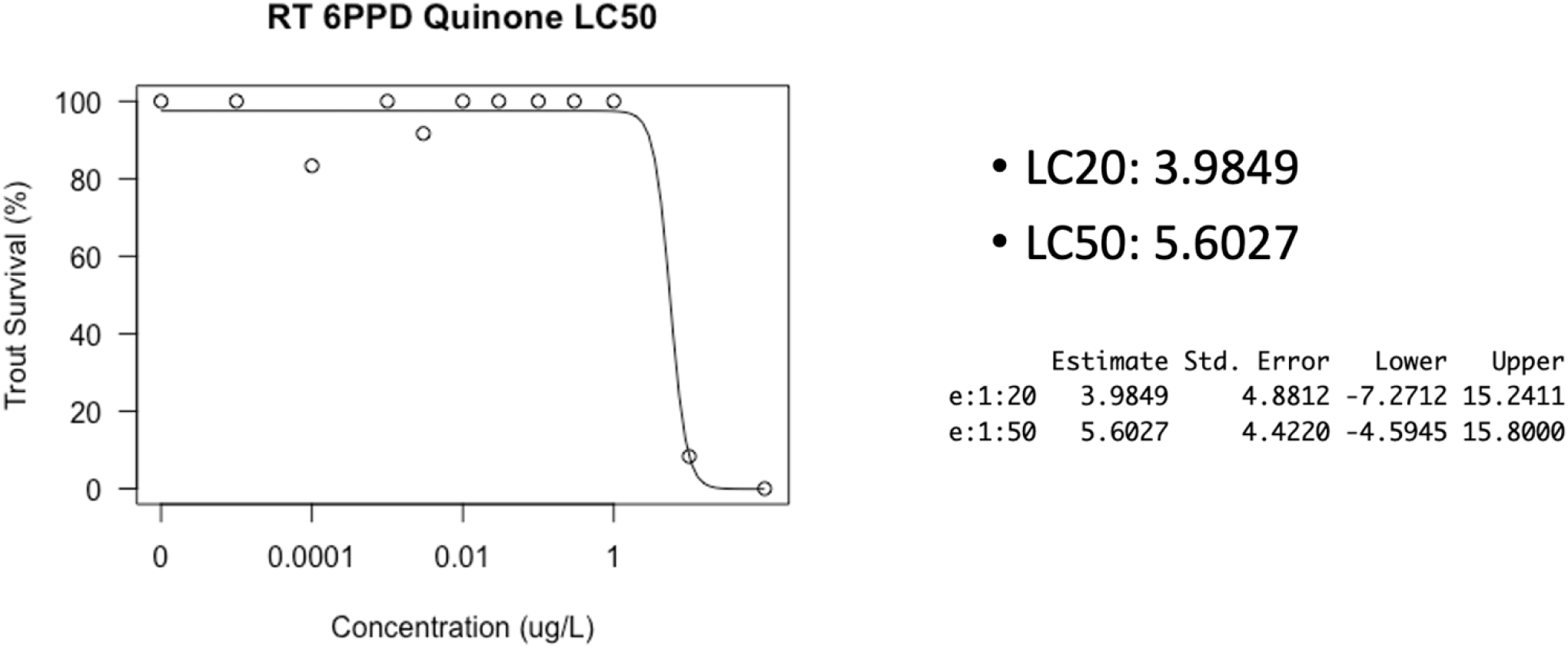

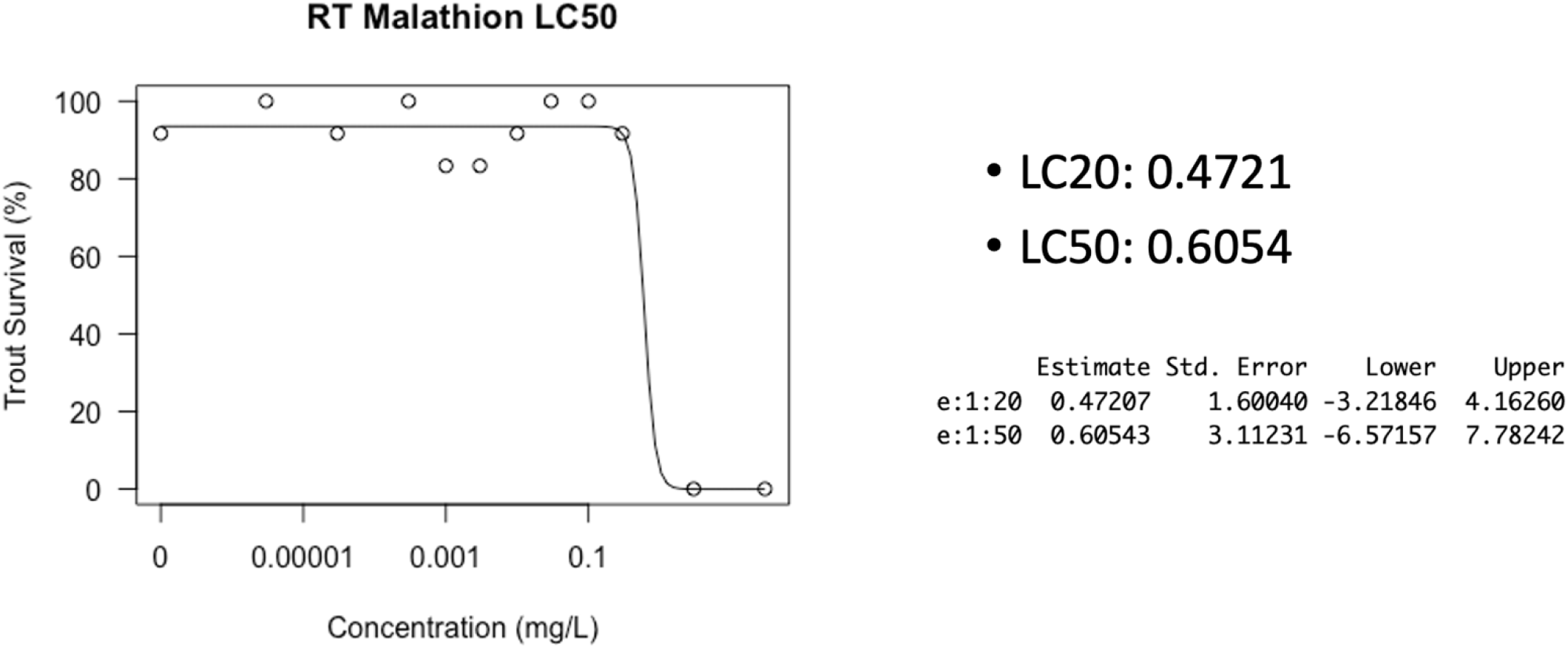

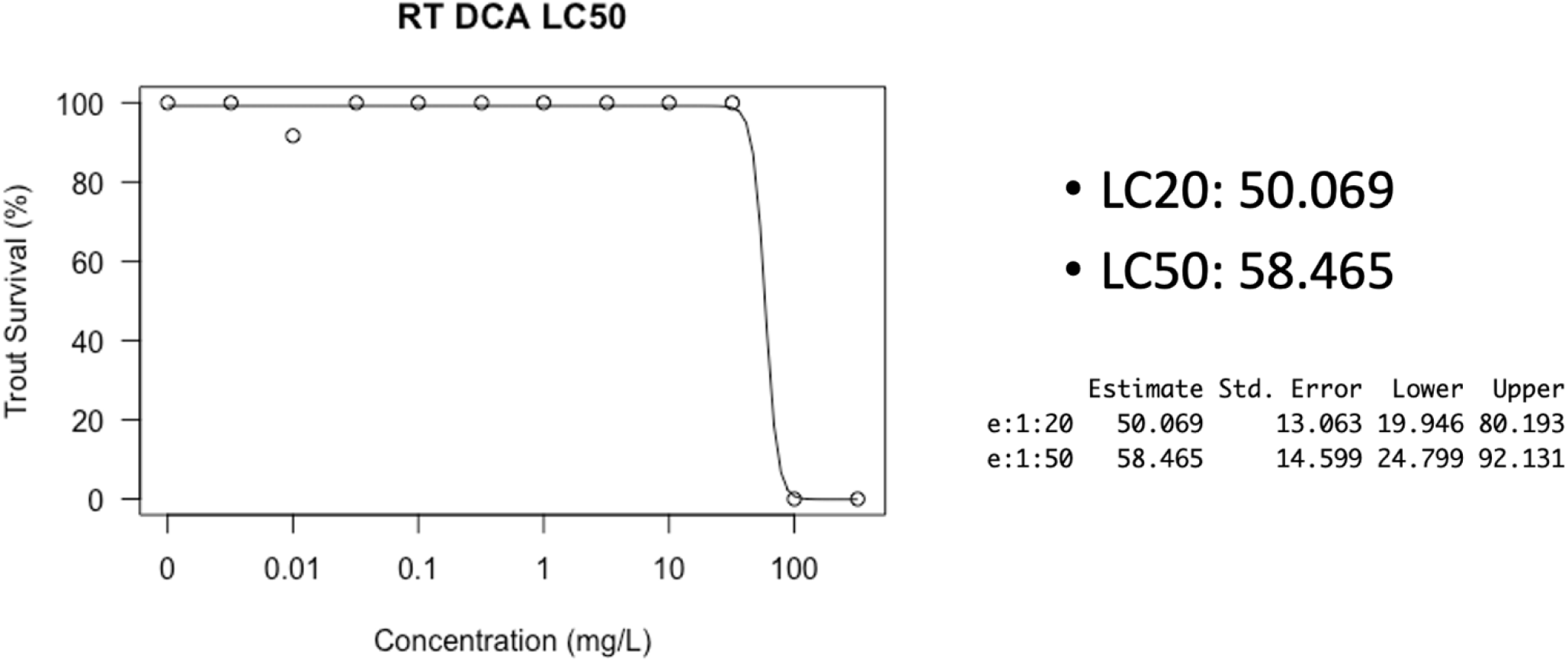

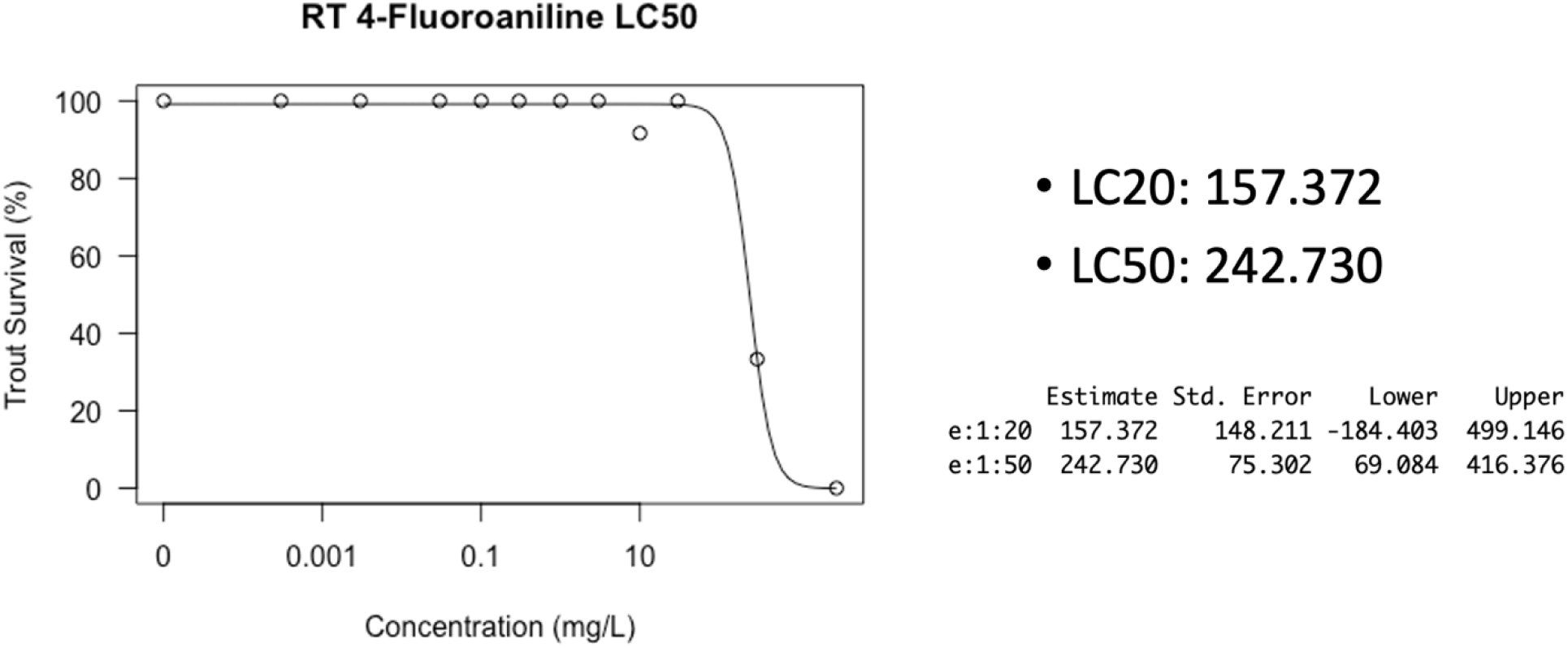

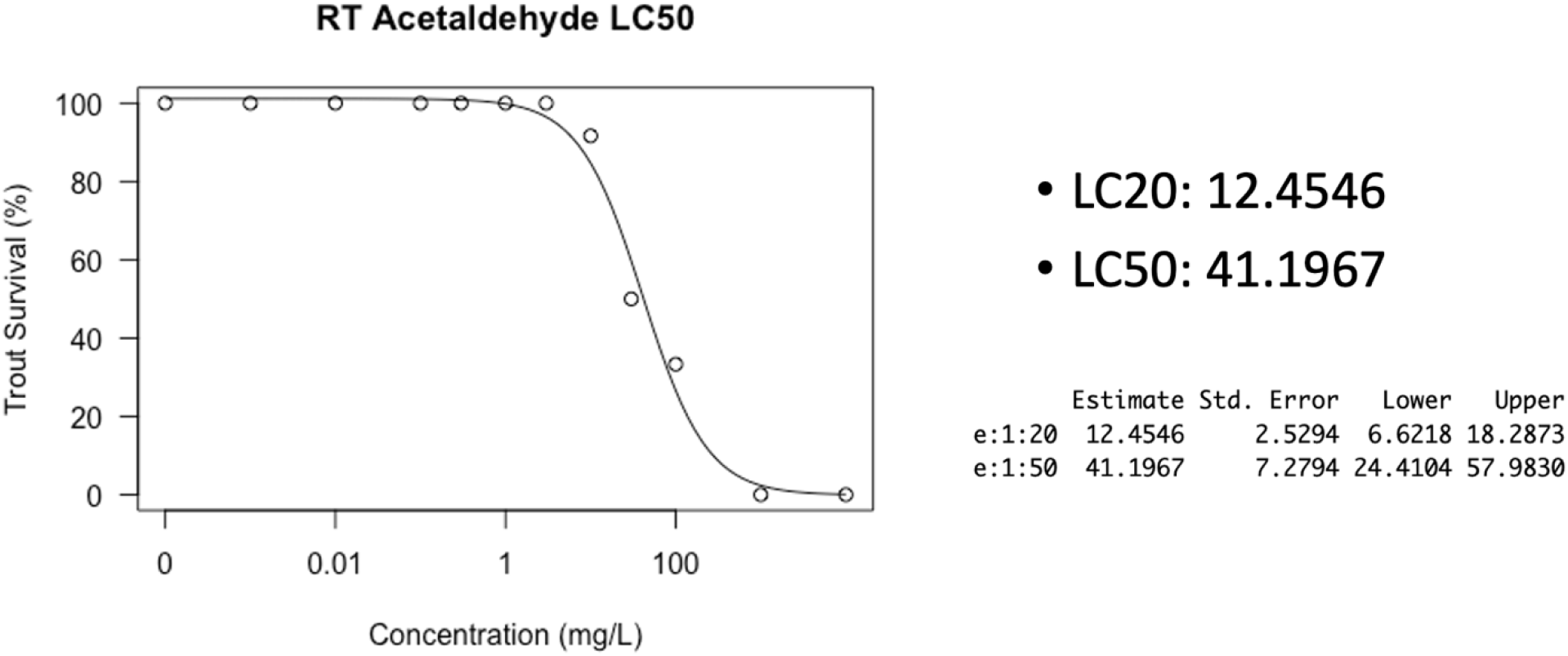

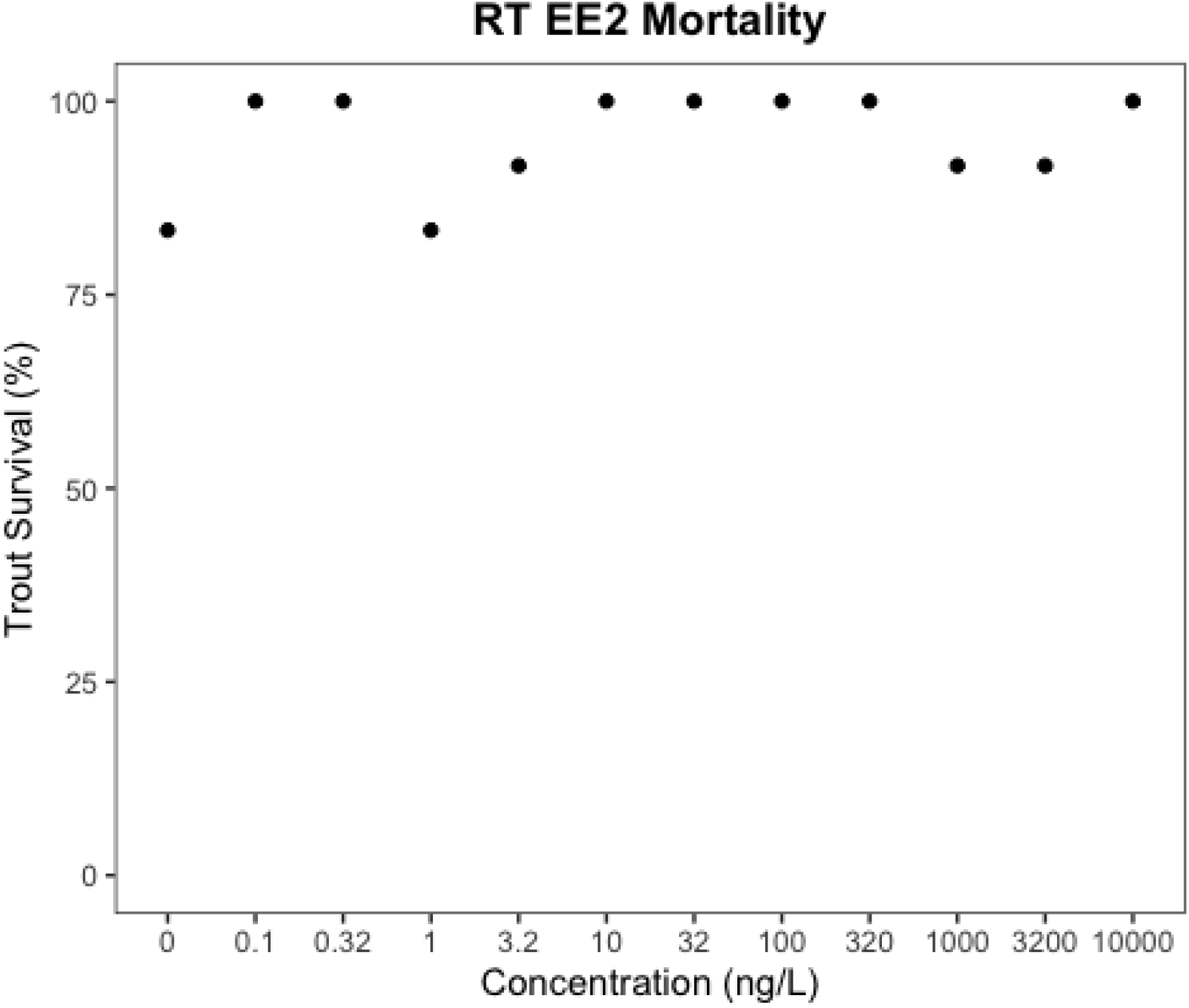

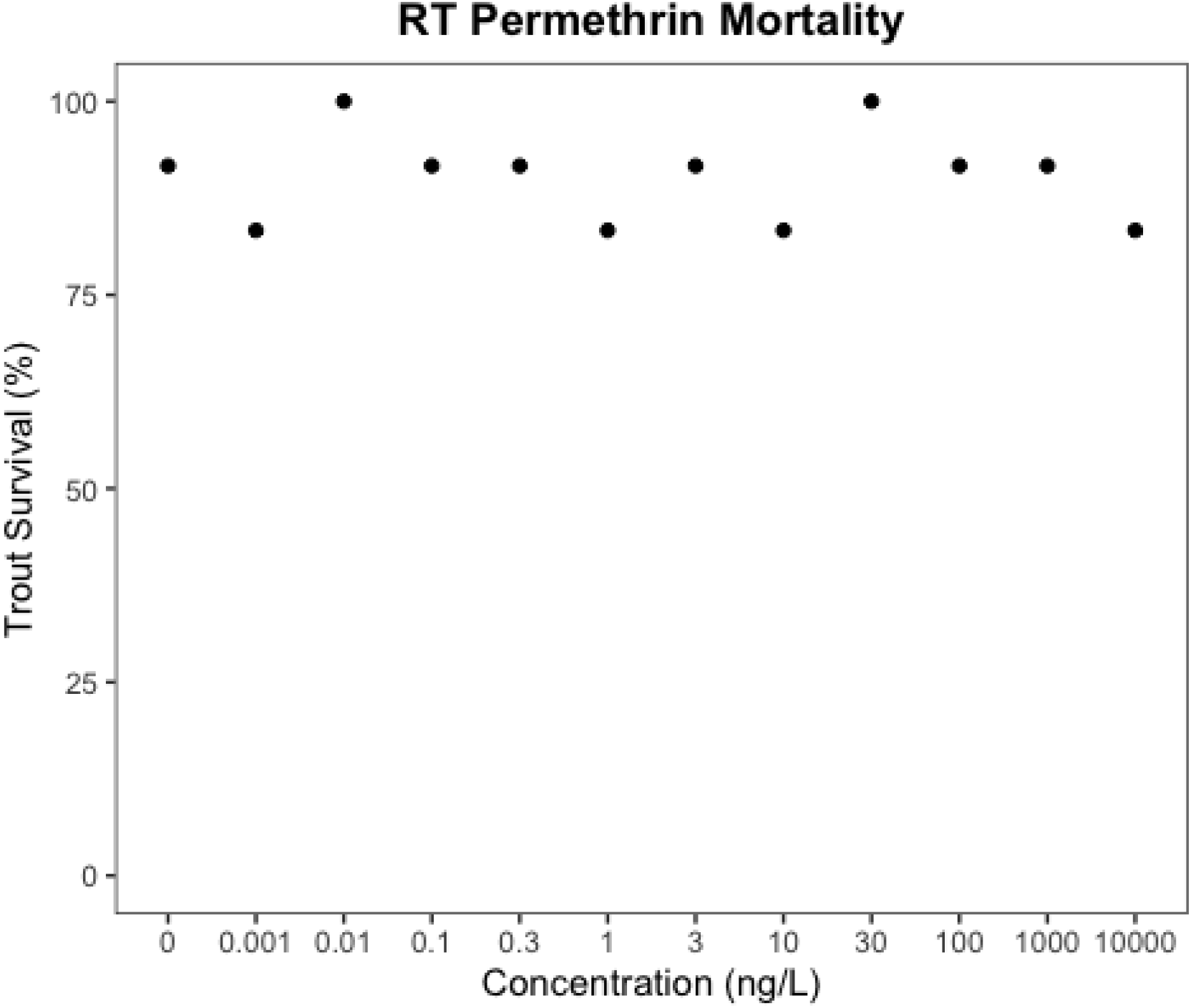

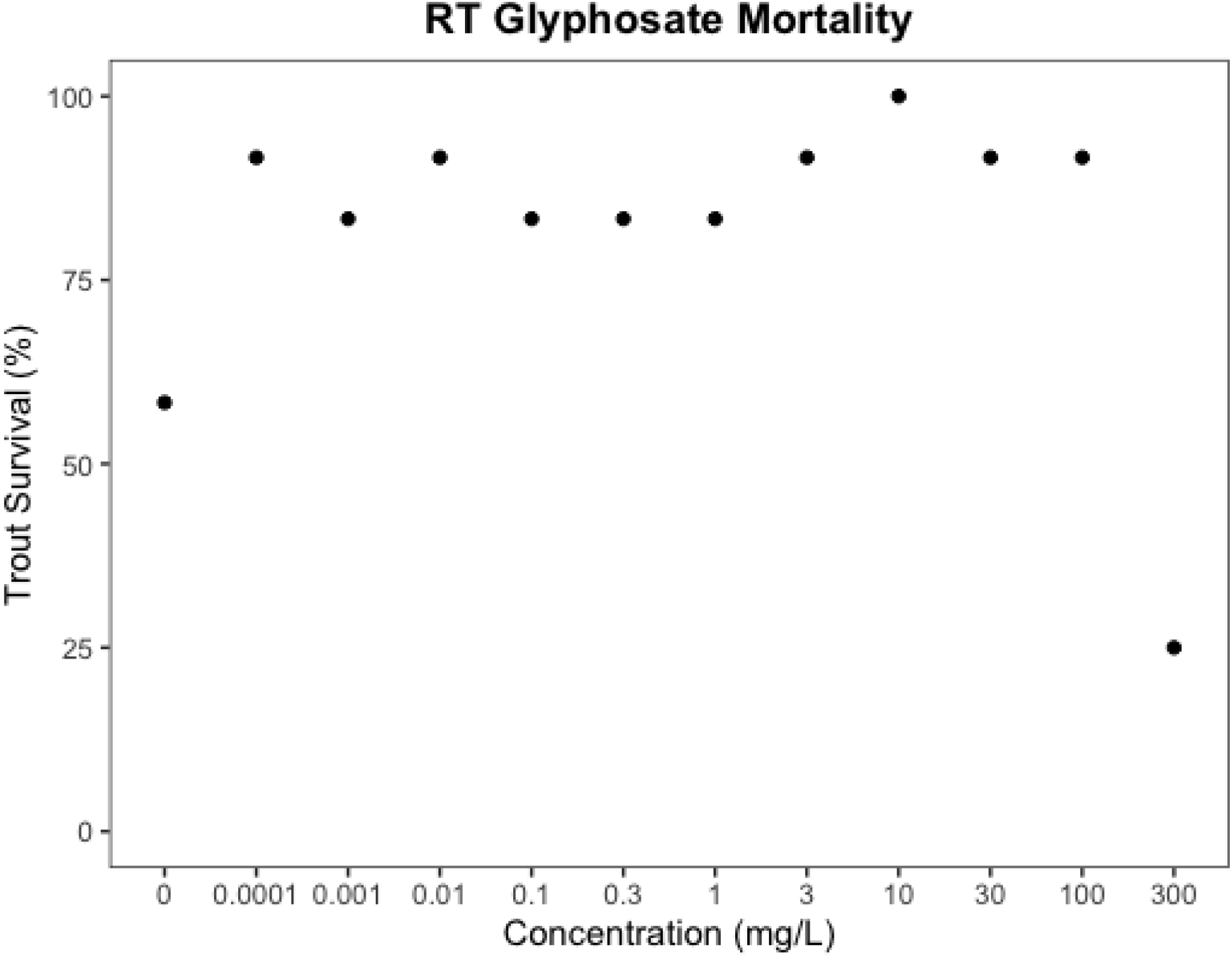

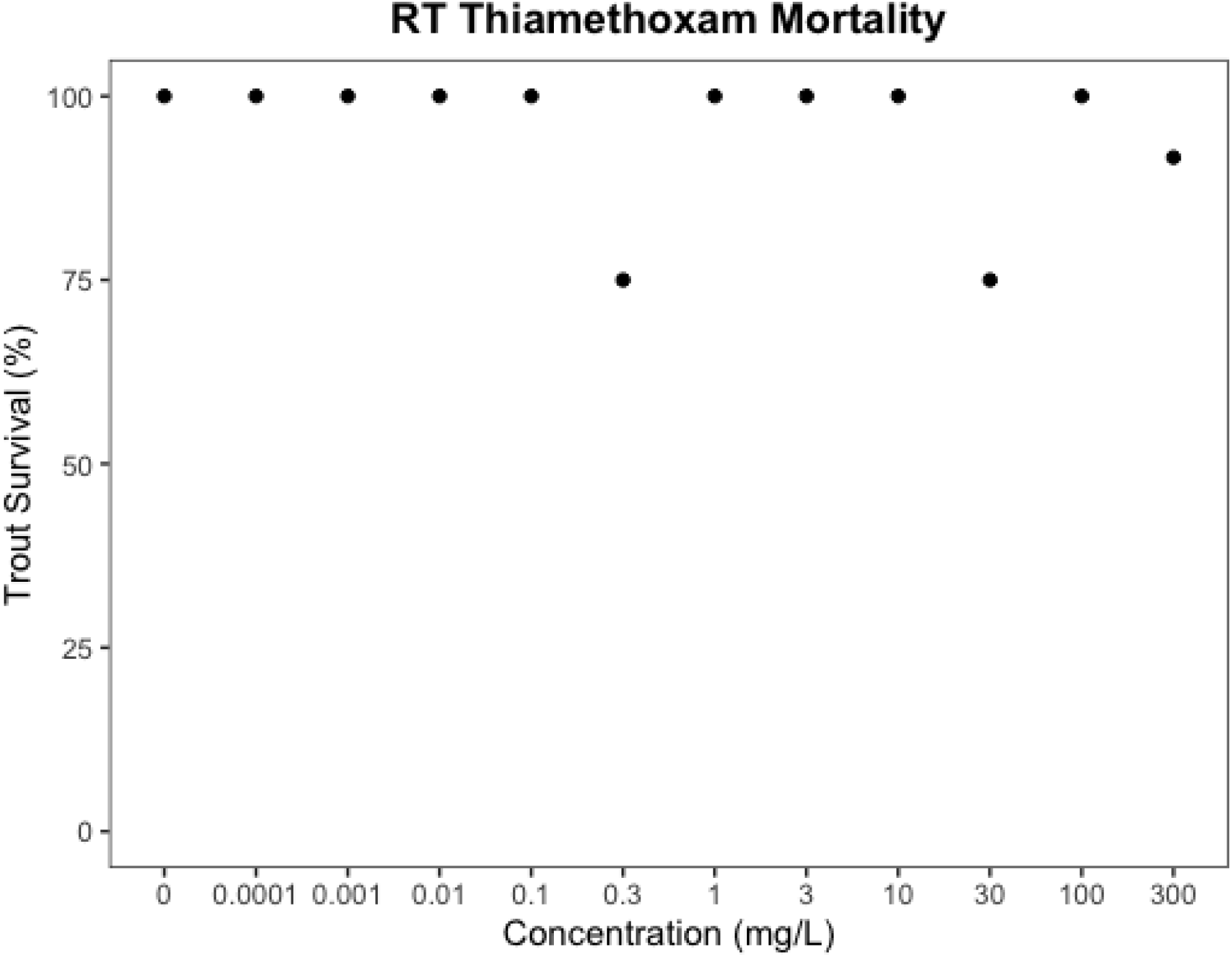

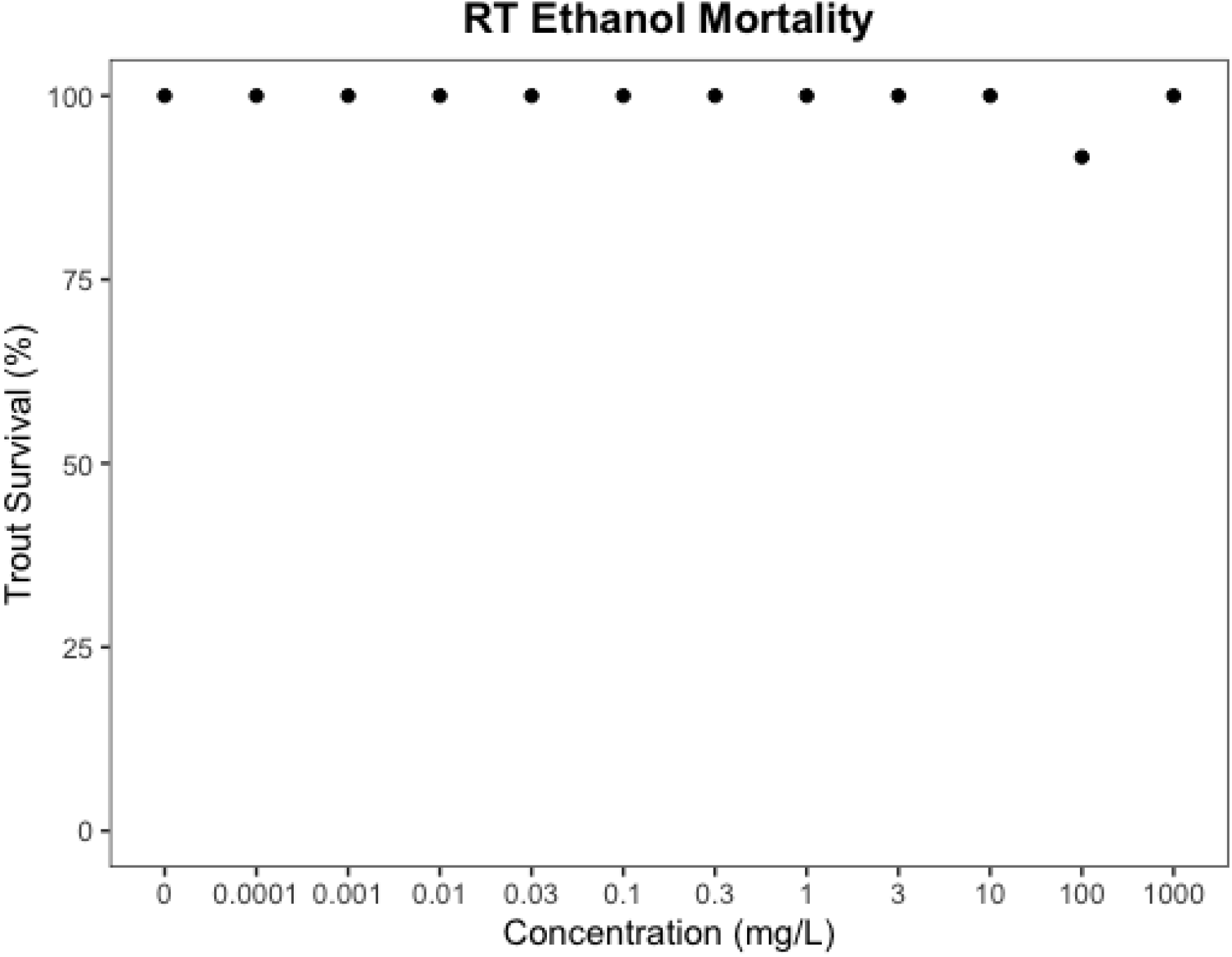

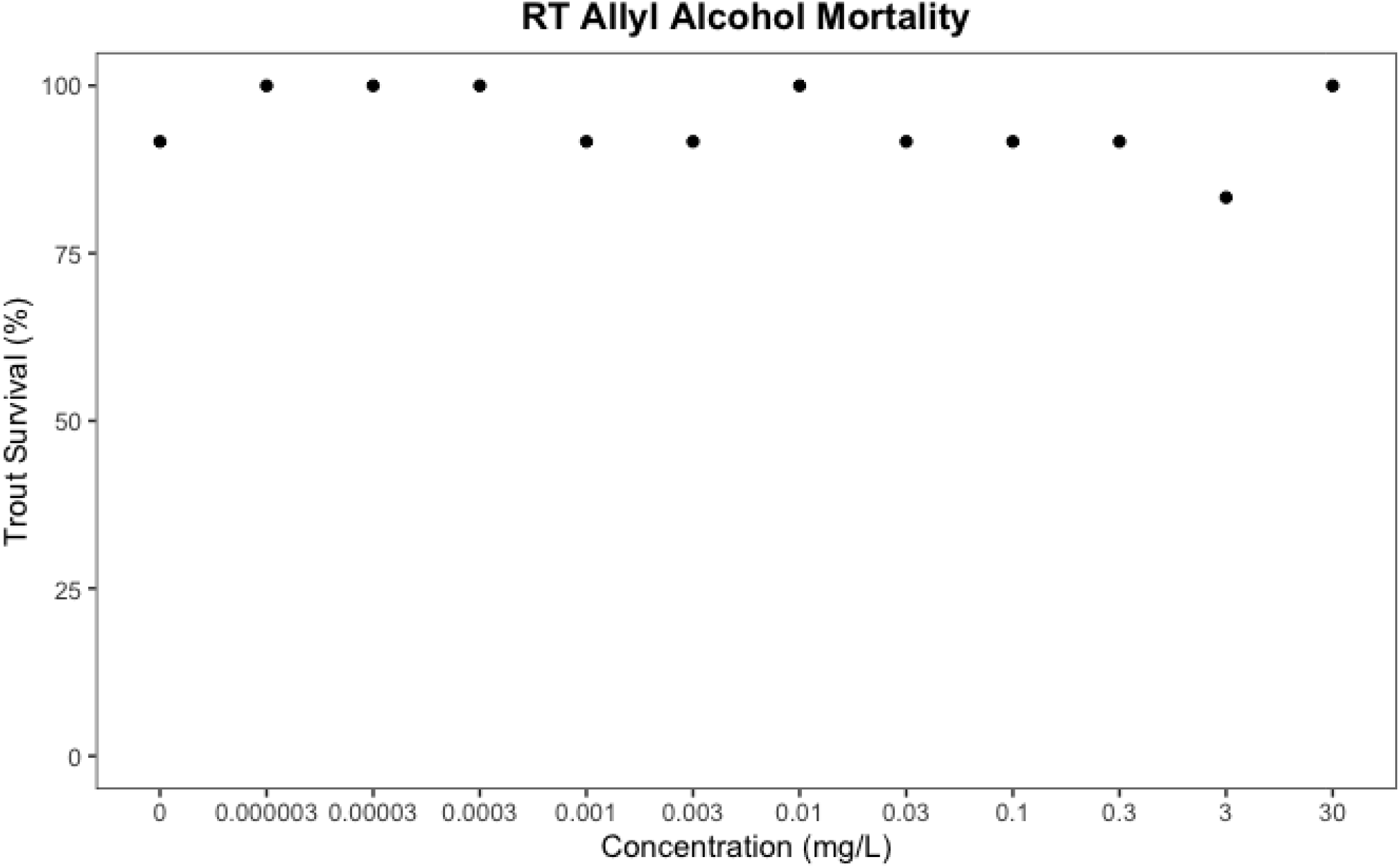

## References Cited

Belanger SE, Balon EK, Rawlings JM. Saltatory ontogeny of fishes and sensitive early life stages for ecotoxicology tests. Aquat Toxicol. 2010 Apr 15;97(2):88–95. doi: 10.1016/j.aquatox.2009.11.020

Bérubé, R., Lefebvre-Raine, M., Gauthier, C., Bourdin, T., Bellot, P., Triffault-Bouchet, G., Langlois, V. S., & Couture, P. (2022). Comparative toxicity of conventional and unconventional oils during rainbow trout (Oncorhynchus mykiss) embryonic development: From molecular to health consequences. Chemosphere, 288, 132521.

Brinkmann, M., Montgomery, D., Selinger, S., Miller, J. G. P., Stock, E., Alcaraz, A. J., Challis, J. K., Weber, L., Janz, D., Hecker, M., & Wiseman, S. (2022). Acute toxicity of the tire rubber-derived chemical 6PPD-quinone to four fishes of commercial, cultural, and ecological importance. Environmental Science & Technology Letters, 9, 333–338. https://doi.org/10.1021/acs.estlett.2c00050

Chen X, He T, Yang X, Gan Y, Qing X, Wang J, Huang Y. Analysis, environmental occurrence, fate and potential toxicity of tire wear compounds 6PPD and 6PPD-quinone. J Hazard Mater. 2023 Jun 15;452:131245. doi: 10.1016/j.jhazmat.2023.131245

Di, S., Liu, Z., Zhao, H., Li, Y., Qi, P., Wang, Z., Xu, H., Jin, Y., & Wang, X. (2022). Chiral perspective evaluations: Enantioselective hydrolysis of 6PPD and 6PPD-quinone in water and enantioselective toxicity to *Gobiocypris rarus* and *Oncorhynchus mykiss*. Environment International, 166, 107374. https://doi.org/10.1016/j.envint.2022.107374

Di Lombo, M. S., Weeks-Santos, S., Clérandeau, C., Triffault-Bouchet, G., Valérie, S. L., Couture, P., & Cachot, J. (2021). Comparative developmental toxicity of conventional oils and diluted bitumen on early life stages of the rainbow trout (Oncorhynchus mykiss). Aquatic Toxicology, 239, 105937.

EChA 2016. New approach methodologies in regulatory science. European Chemicals Agency https://echa.europa.eu/documents/10162/21838212/scientific_ws_proceedings_en.pdf/a2087434-0407-4705-9057-95d9c2c2cc57.

Embry MR, Belanger SE, Braunbeck TA, Galay-Burgos M, Halder M, Hinton DE, Léonard MA, Lillicrap A, Norberg-King T, Whale G. The fish embryo toxicity test as an animal alternative method in hazard and risk assessment and scientific research. Aquat Toxicol. 2010 Apr 15;97(2):79–87. doi: 10.1016/j.aquatox.2009.12.008. Epub 2009 Dec 16. PMID: 20061034.

Halder M, Léonard M, Iguchi T, Oris JT, Ryder K, Belanger SE, Braunbeck TA, Embry MR, Whale G, Norberg-King T, Lillicrap A. Regulatory aspects on the use of fish embryos in environmental toxicology. Integr Environ Assess Manag. 2010 Jul;6(3):484–91. doi: 10.1002/ieam.48. PMID: 20821708.

Hoyberghs J, Bars C, Ayuso M, Van Ginneken C, Foubert K, Van Cruchten S. DMSO Concentrations up to 1% are Safe to be Used in the Zebrafish Embryo Developmental Toxicity Assay. Front Toxicol. 2021 Dec 21;3:804033. doi: 10.3389/ftox.2021.804033

Johnson KJ, Auerbach SS, Stevens T, Barton-Maclaren TS, Costa E, Currie RA, Dalmas Wilk D, Haq S, Rager JE, Reardon AJF, Wehmas L, Williams A, O’Brien J, Yauk C, LaRocca JL, Pettit S. A Transformative Vision for an Omics-Based Regulatory Chemical Testing Paradigm. Toxicol Sci. 2022 Nov 23;190(2):127–132. doi: 10.1093/toxsci/kfac097

Morash MG, Kirzinger MW, Achenbach JC, Venkatachalam AB, Cooper JP, Ratzlaff DE, Woodland CLA, Ellis LD. The contribution of larval zebrafish transcriptomics to chemical risk assessment. Regul Toxicol Pharmacol. 2023 Feb;138:105336. doi: 10.1016/j.yrtph.2023.105336.

National Toxicology Program NTP Research Report on National Toxicology Program Approach to Genomic Dose-Response Modeling. NTP RR 5. Research Triangle Park. NC National Toxicology Program. 2018;5:1–44.

Pagé-Larivière F, Crump D, O’Brien JM. Transcriptomic points-of-departure from short-term exposure studies are protective of chronic effects for fish exposed to estrogenic chemicals. Toxicol Appl Pharmacol. 2019 Sep 1;378:114634. doi: 10.1016/j.taap.2019.114634

United Nations. 2015. Globally harmonized system of classification and labelling of chemicals (GHS). Sixth Revised Edition. ST/SG/AC.10/30/Rev.6. United Nations, New York, NY. Part 4 – Environmental Hazards

U.S. Environmental Protection Agency (2018) Strategic plan to promote the development and implementation of alternative test methods within the TSCA program. https://www.epa.gov/sites/production/files/2018-06/documents/epa_alt_strat_plan_6-20-18_clean_final.pdf

van der Zalm AJ, Barroso J, Browne P, Casey W, Gordon J, Henry TR, Kleinstreuer NC, Lowit AB, Perron M, Clippinger AJ. A framework for establishing scientific confidence in new approach methodologies. Arch Toxicol. 2022 Nov;96(11):2865–2879. doi: 10.1007/s00204-022-03365-4

Villeneuve DL, Le M, Hazemi M, Biales A, Bencic DC, Bush K, Flick R, Martinson J, Morshead M, Rodriguez KS, Vitense K, Flynn K. Pilot testing and optimization of a larval fathead minnow high throughput transcriptomics assay. Curr Res Toxicol. 2022 Dec 22;4:100099. doi: 10.1016/j.crtox.2022.100099

Weeks Santos, S., Gonzalez, P., Cormier, B., Mazzella, N., Moreira, A., Clérandeau, C., Morin, B., & Cachot, J. (2021). Subchronic Exposure to Environmental Concentrations of Chlorpyrifos Affects Swimming Activity of Rainbow Trout Larvae. Environmental Toxicology and Chemistry, 40(11), 3092–3102.

